## Supplementary Figures S1-S17, Table S1, Notes S1-S5 for "Joint, multifaceted genomic analysis enables diagnosis of diverse, ultra-rare monogenic presentations"

Figure S13. Enrichment of correct diagnoses in RaMeDiES-IND's patient–gene ranking.14

### Figures

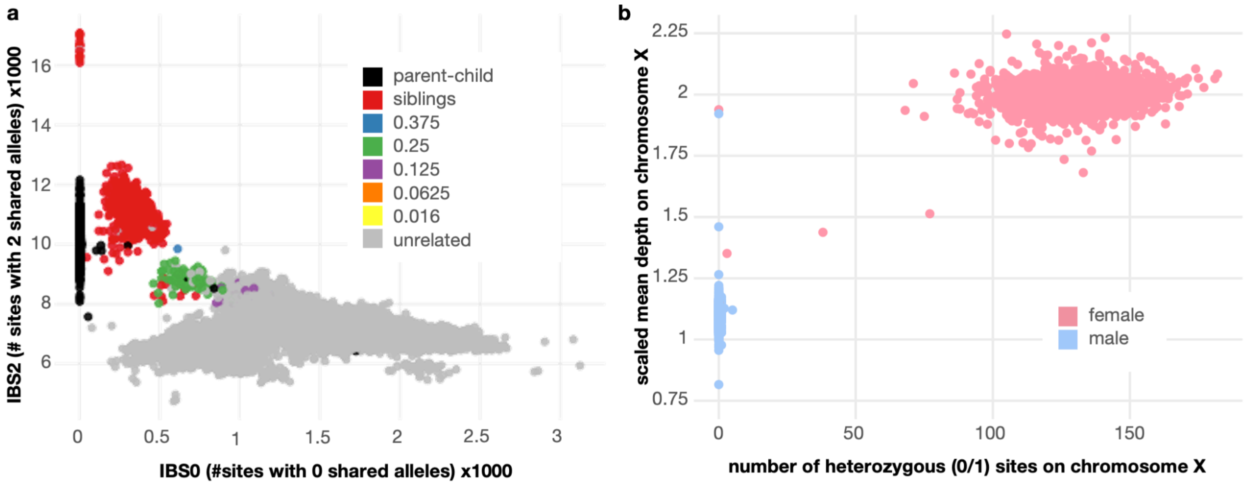

**Figure S1. UDN cohort pairwise relatedness and sex quality control checks.**
**(a)** Between-sample pairwise relatedness computed using Somalier, which operates on a predefined, informative subset of genomic positions.[1] Unrelated individuals are expected to have a nonzero number of genomic positions with no shared alleles (IBS0), whereas parent–child pairs should have at least one shared allele at every genomic site. **(b)** Within-sample metrics for checking the sex of samples; females (XX, pink) are expected to have heterozygous sites on chromosome X and a scaled mean depth (calculated as two times the read depth on chromosome X / mean read depth across the genome) of about 2.

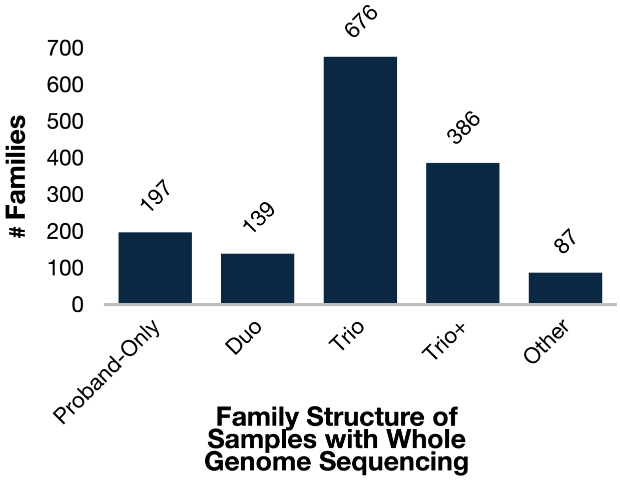

**Figure S2. Family structure of sequenced individuals.**
Duo cases are not necessarily always parent–child relationships; some sibling pairs are recorded as duos. Trios (and trio+) families require at least one child with both parents sequenced. Families with three sequenced individuals who do not share this relationship pattern are included in the “other” category.

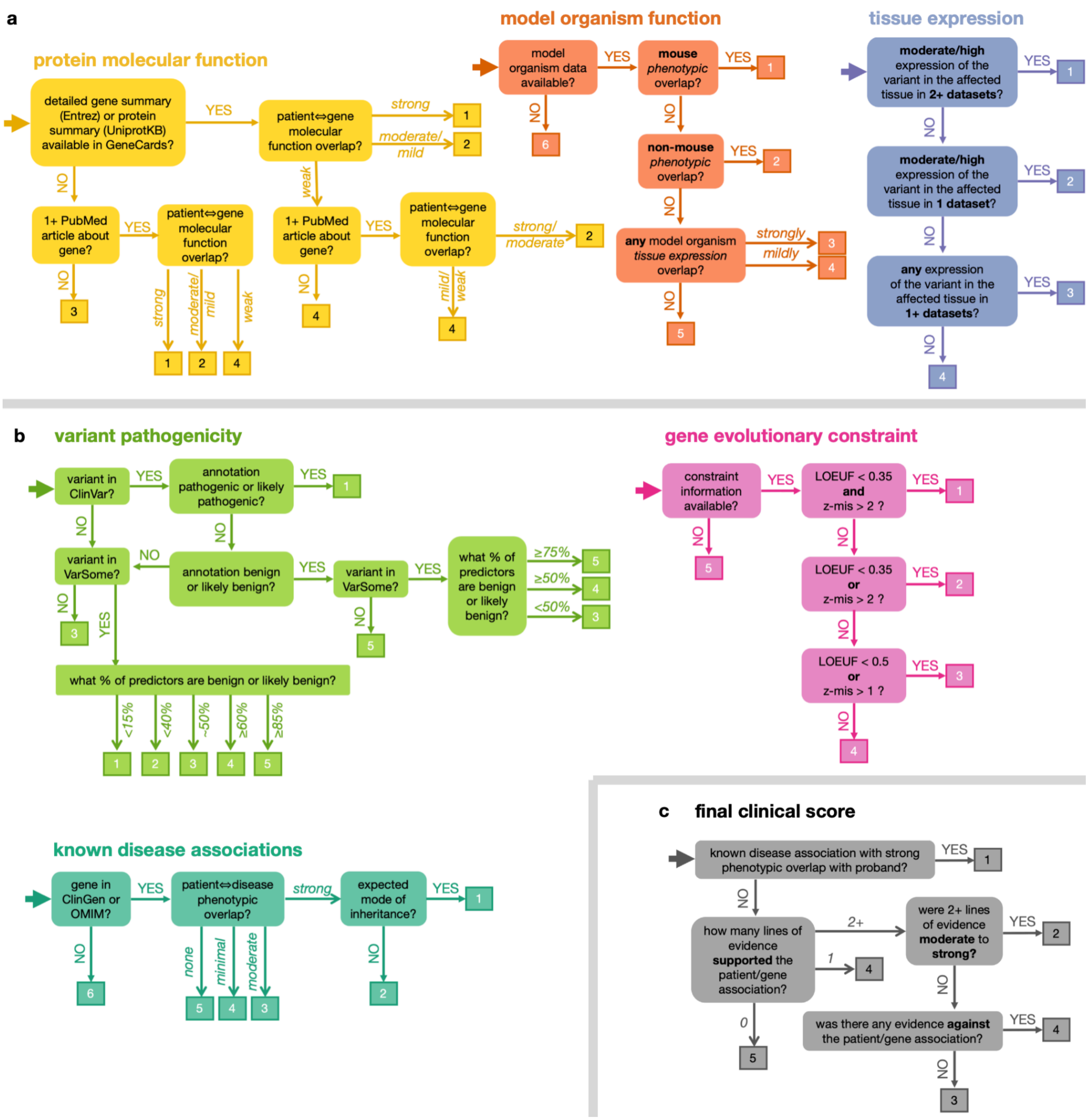

**Figure S3. Hierarchical decision trees for clinically assessing candidates.**

**(a)** Three categories of evidence used in the clinical evaluation protocol for assessing gene-disease fit for each participant phenotype that were not used in any statistical candidate prioritization methods. **(b)** Three categories of evidence used where clinicians are cognizant of how evidence assessed may be reflective of or correlated with features that were used in the statistical candidate prioritization. **(c)** Final clinical score assigned based on the six categories of evidence.

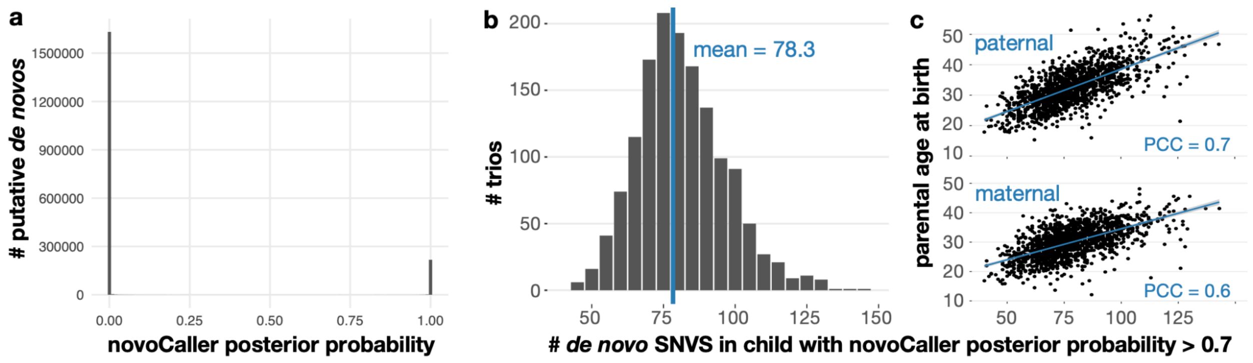

**Figure S4. NovoCaller selects for confident *de novo* variants.**

Putative *de novo* variants were selected in each trio as described in Methods. **(a)** Posterior probabilities assigned by novoCaller are bimodal; 40.4% of putative *de novos* were assigned a posterior probability of 0 and are excluded from the histogram. **(b)** Counts of *de novo* SNVs per child in each trio uncorrected by parental ages. **(c)** Counts of *de novo* SNVs show expected correlation with paternal (top) and maternal (bottom) age.

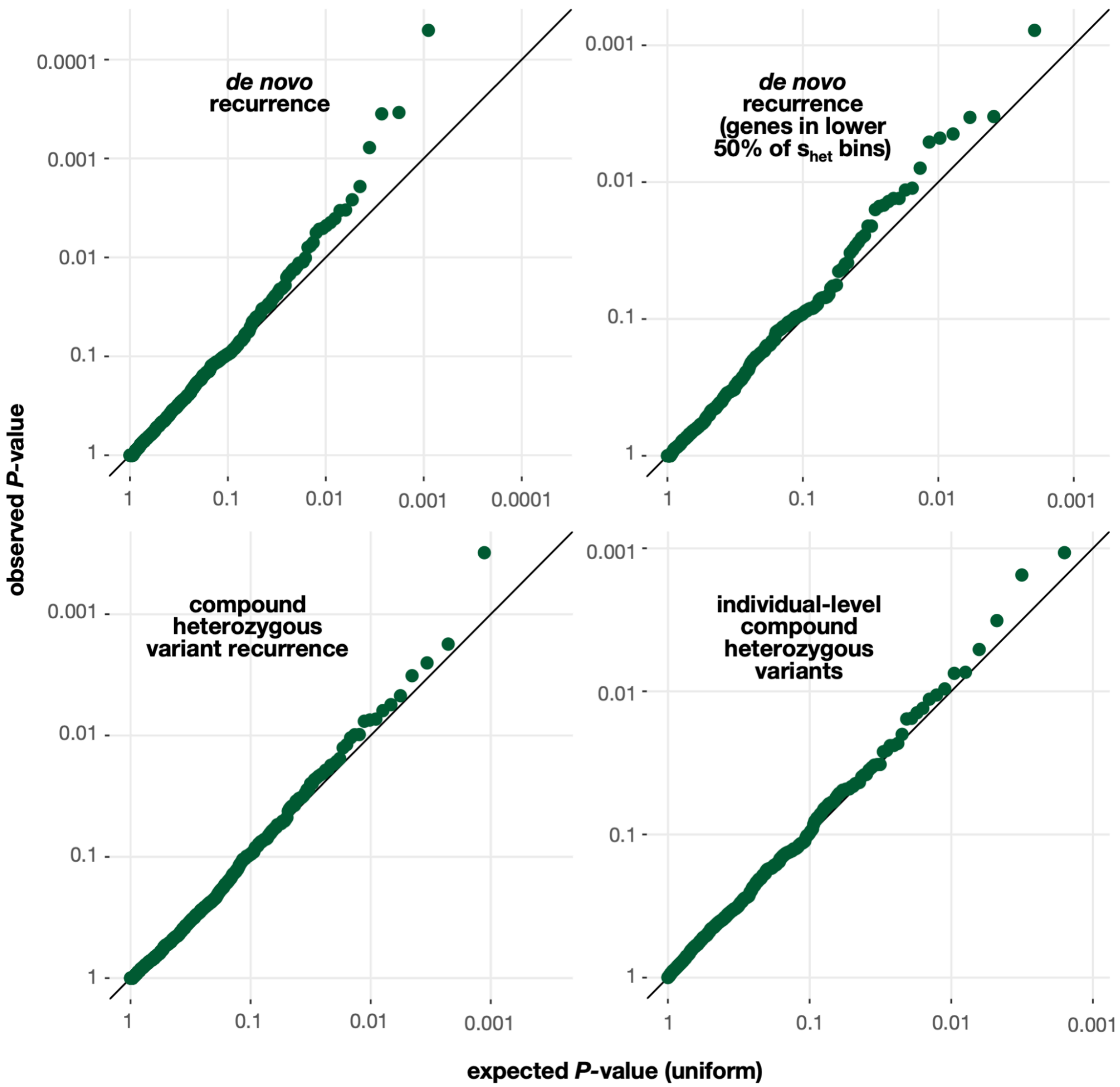

**Figure S5. QQ-plots for closed-form *de novo* and compound heterozygous statistics.** Top row depicts QQ-plots for *de novo* recurrence. The full curve is shown on the left and the subset of genes falling into the lower half of $s_{het}$ bins are shown on the right. The bottom row depicts QQ-plots for compound heterozygous recurrence (left) and individual-level (right) statistics. Because all of our closed-form statistics $y$ and $Pr(y|K)$ are not defined for $K$=0 and therefore not expected to be uniformly-distributed (see Methods), the $P$-values plotted here are $Pr(y|K\geq1) = Pr(y)/Pr(K\geq1)$.

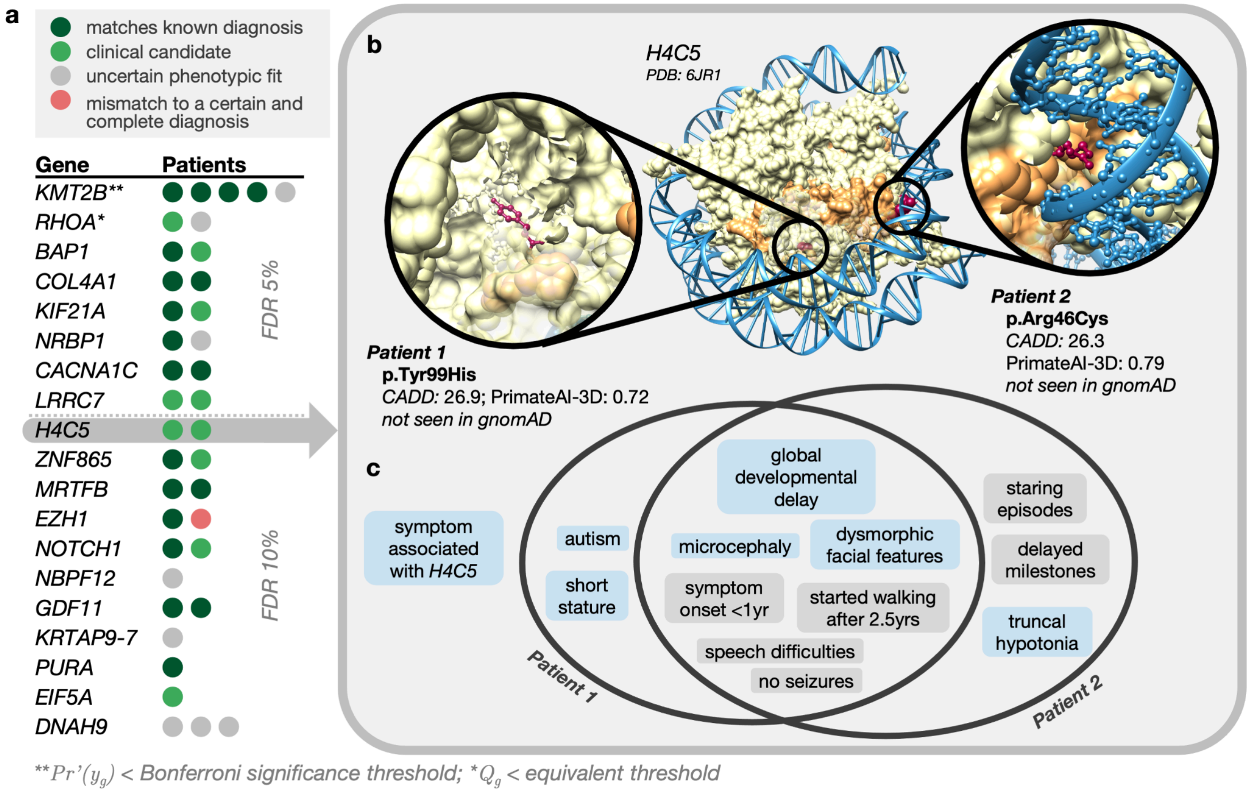

**Figure S6. Genes recurrently hit with exonic variants highlight *H4C5*.**
**(a)** Gene findings at FDR 5% and 10% when running RaMeDiES-DN on all exonic variants (including indels and nonsense variants) using four deleteriousness predictors (AlphaMissense, PrimateAI-3D, CADD and REVEL). **(b)** Human histone protein structure (PDB ID: 6JR1) where chains B and F, sequence matches to the *H4C5* gene, are highlighted in orange. Reference alleles for missense *de novo* variants observed across two UDN patients (red) are shown in zoomed-in circles. **(c)** Overlap of phenotype terms annotated to each patient. Phenotypes associated with histone genes or *H4C5* specifically are highlighted in blue, and phenotypes not yet associated with histone genes are highlighted in gray.

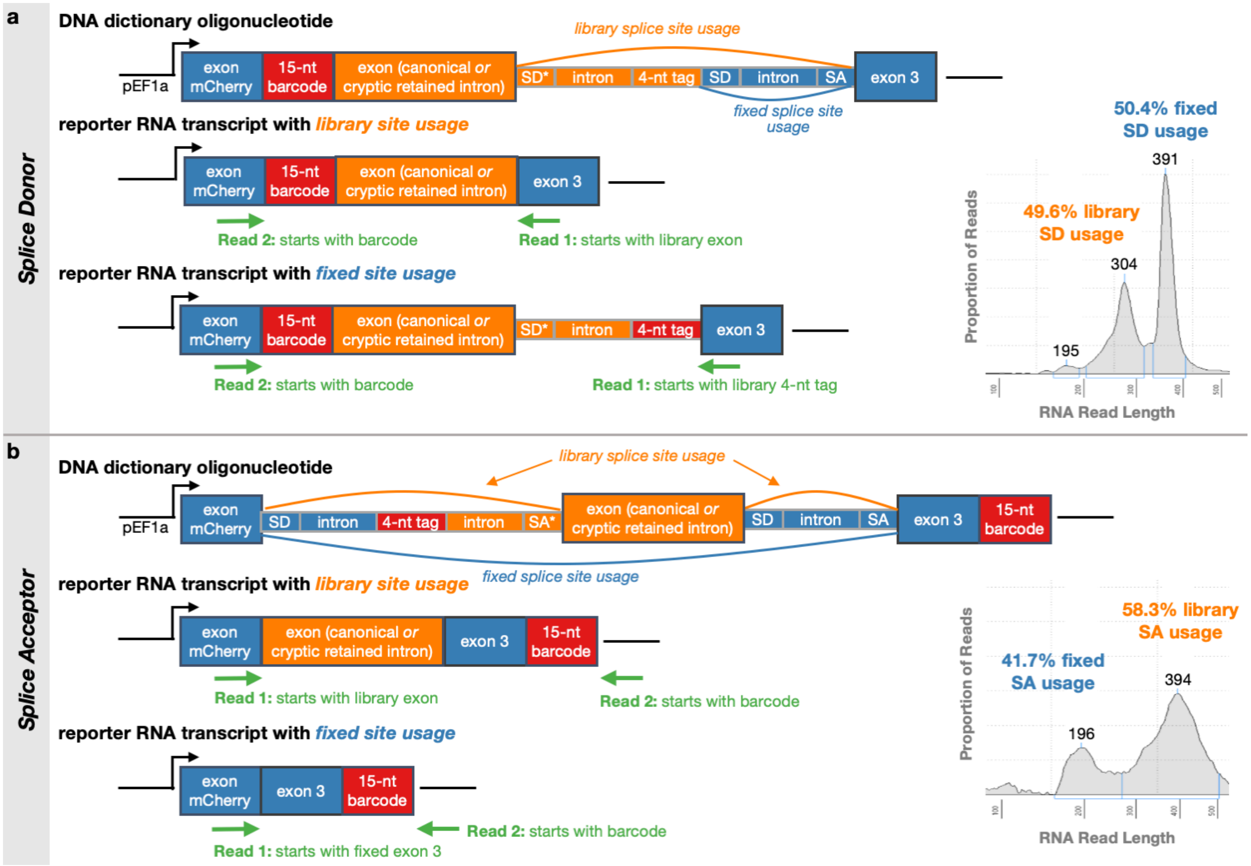

**Figure S7. Massively Parallel Splicing Reporter Assay (MPSA) design**

Oligonucleotides containing a 155-nucleotide section of genomic DNA from a UDN patient, called a 'library' (orange) were inserted into lentiviral plasmids (pEF1a). Exons are represented as tall boxes and introns as short boxes in the DNA depictions. These genomic DNA library segments were uniquely tagged with barcodes detectable in DNA- or RNA-sequencing (red). Libraries contained either canonical splice sites that a variant was predicted to cause the loss of, or cryptic splice sites that a variant was predicted to cause the gain of (denoted with an '*'). Fixed exonic and intronic sequence with fixed splice sites surround the inserted library (blue). Expected paired RNA-sequencing reads are depicted in green. **(a)** Usage of a library splice donor site versus the fixed splice donor site can be detected in reporter RNA transcripts. Inset shows the proportions of RNA reads of different lengths produced through Tapestation, an automated electrophoresis system for sizing and quantifying nucleic acid sequence reads. **(b)** Similarly, usage of a library splice acceptor site versus a fixed splice acceptor site can be detected in reporter RNA transcripts.

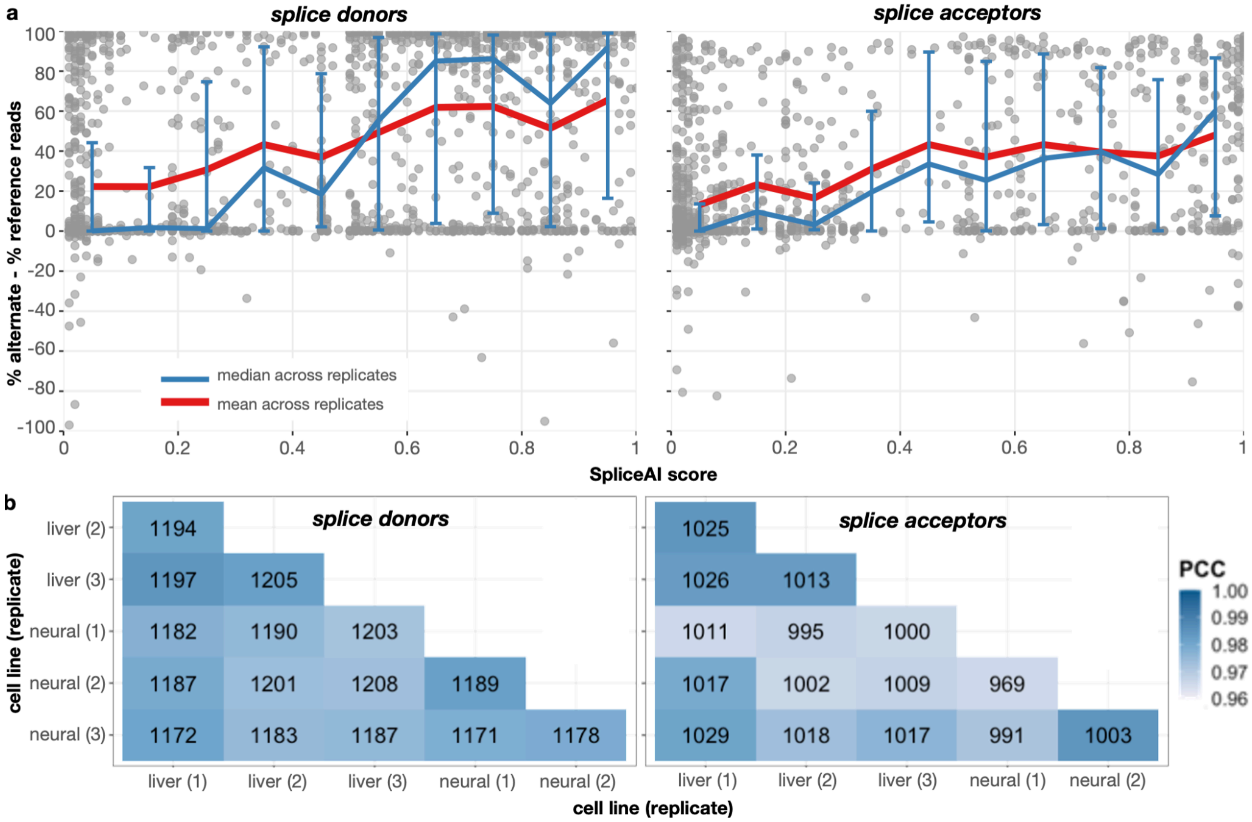

**Figure S8. MPSA validation rates by SpliceAI scores and across replicates.**
A "validation rate" is computed for every variant tested in the MPSA as described in Methods. **(a)** Each point represents the MPSA validation rate for a specific variant that was tested. Blue lines correspond to the median validation rate for all variants within each SpliceAI score decile across biological replicates; error bars depict the interquartile range of validation rates for variants within each decile across replicates. Red lines correspond to the mean validation rate for all variants within each SpliceAI score decile. **(b)** We computed all-against-all Pearson correlations between MPSA validation rates corresponding to the same alternate/reference pairs across experiments. The numbers in the center of each cell correspond to the number of alternate/reference pairs with at least 10 supporting reads each in both experiments being compared. The number in parentheses in the axis labels indicates the biological replicate number. Liver cells are HepG2 and neural-like cells are SK-N-SH.

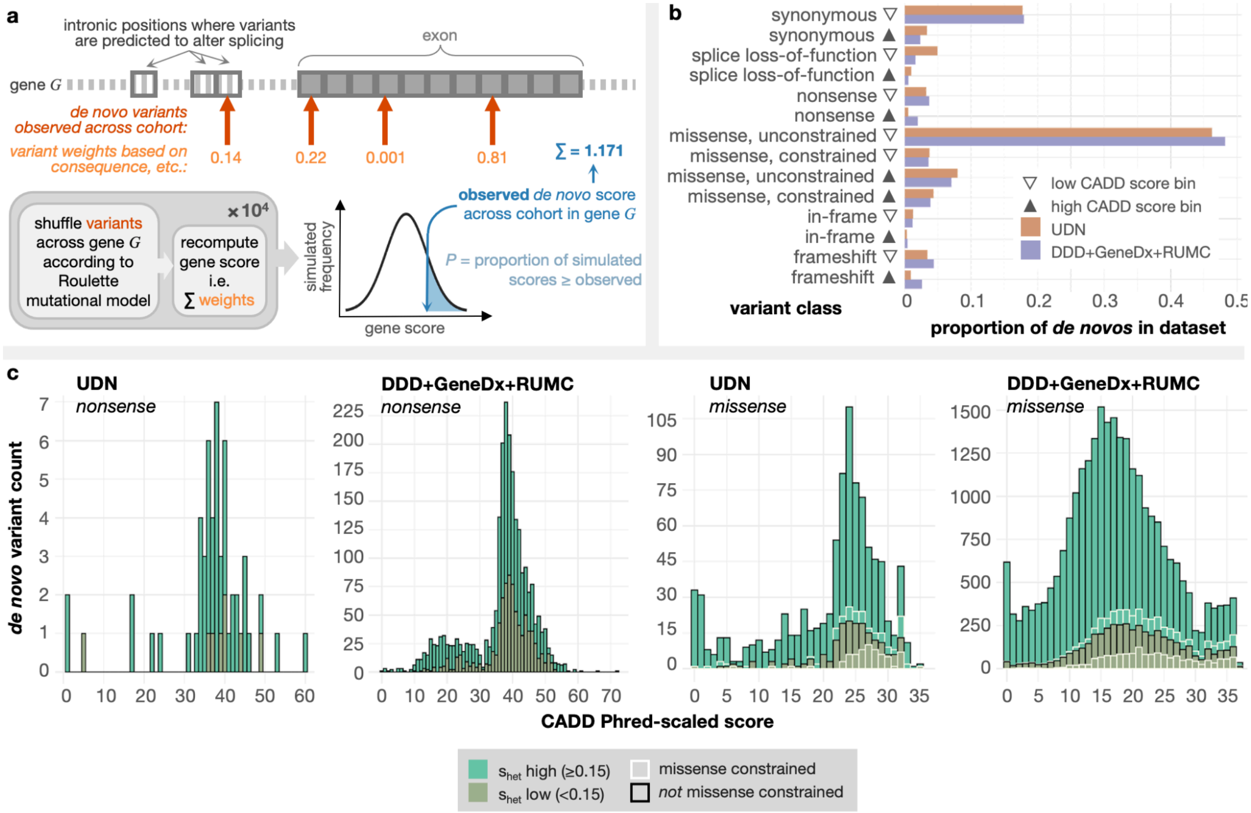

**Figure S9. DeNovoWEST modifications and variant distributions used to calculate weights.**

**(a)** DeNovoWEST incorporates a weighted enrichment test in its loss-of-function (all variants) and gain-of-function (missense variants only) models. We modified this portion of the software to incorporate deeply-intronic variants with a predicted splice-altering impact and replaced the trinucleotide mutation rate model with Roulette as described in Methods.[2] Weights assigned to each variant class were recomputed using the variant type distributions across the UDN and existing DDD+GeneDx+RUMC datasets together. **(b)** Distributions of variants across each variant type bin are largely similar between the UDN and the DDD+GeneDx+RUMC datasets. **(c)** In the nonsense and missense variant type categories, CADD score distribution is similar. We note that the full continuous scale of CADD scores is not used in DeNovoWEST, and variants are binned according to their CADD score instead.

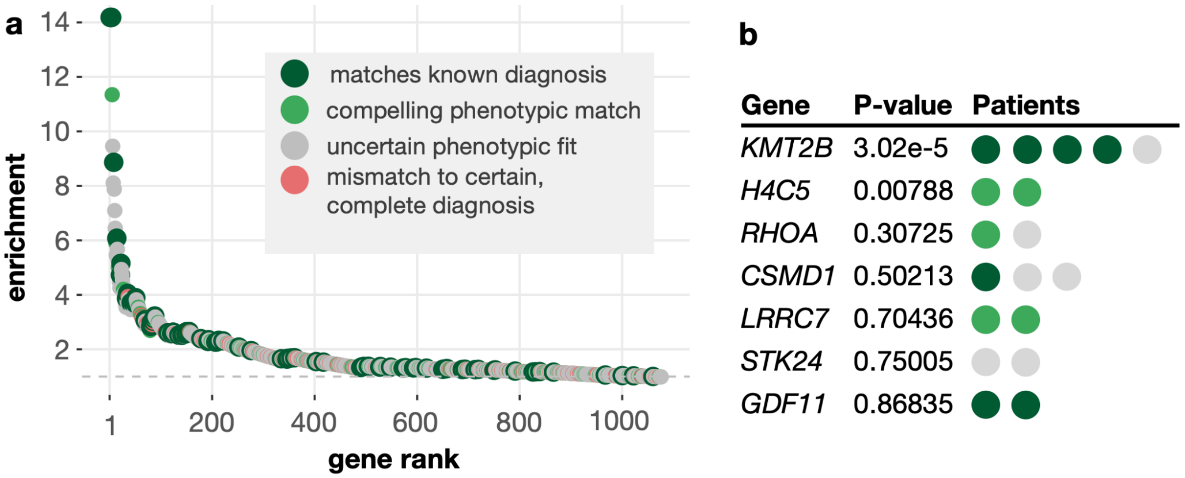

**Figure S10. DeNovoWEST results are enriched for correct diagnoses.**
Genes with observed *de novo* variants across the UDN cohort were evaluated by DeNovoWEST for significant *de novo* recurrence. **(a)** Enrichment for correct diagnoses (green) is calculated as the proportion of correct diagnoses across the top *k* genes divided by the total proportion of correct diagnoses across all genes with observed *de novo* variants. Light green corresponds to an annotated candidate gene. **(b)** Top genes found by DeNovoWEST with Bonferroni-corrected *P*-value. Here, light green corresponds to either a previously annotated candidate gene or a gene that our clinical team found to be a compelling phenotypic match.

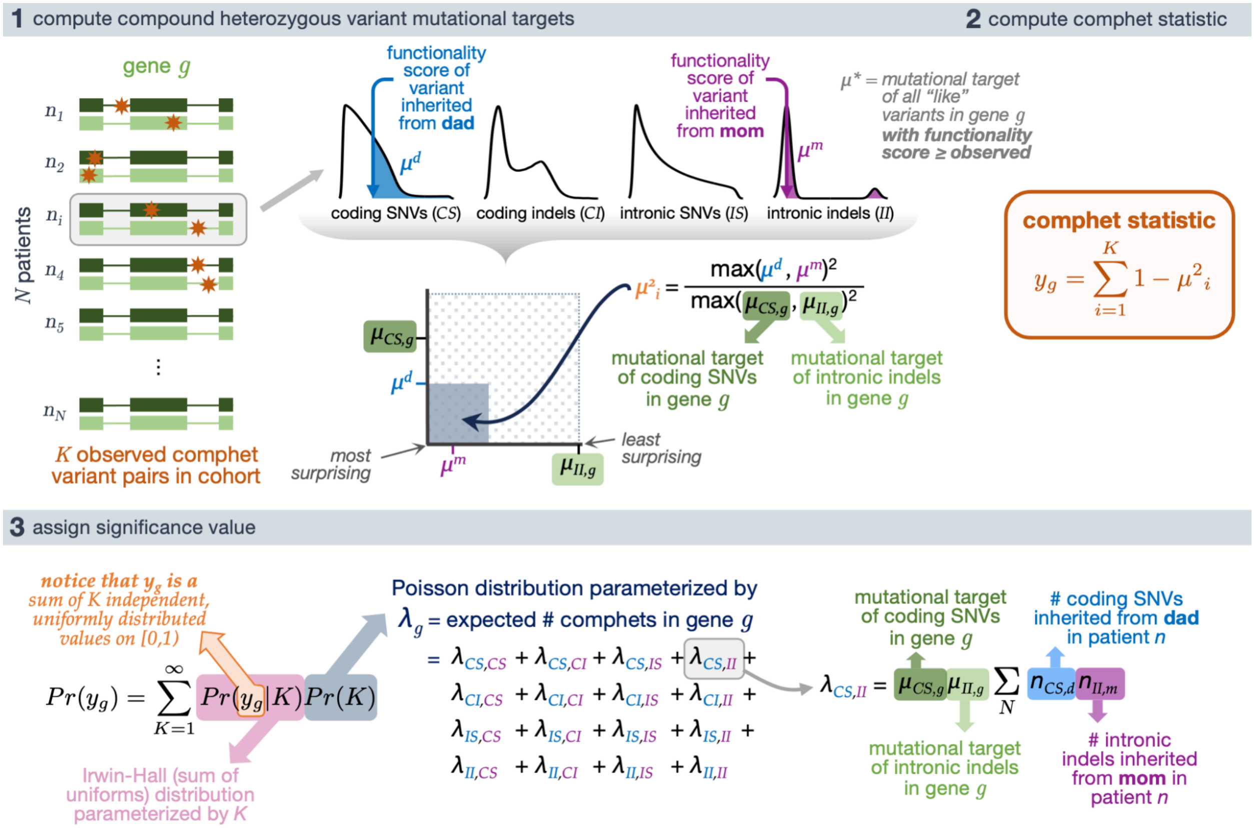

**Figure S11. Schematic of cohort-level compound heterozygous variant recurrence**
Schematic of analytical test for the recurrence of compound heterozygous variant pairs (comphets) that considers distal splice-altering and exonic SNV and indel variants, their variant functionality scores, and a genome-wide mutation rate model Roulette. "Like" variants refer to those of the same variant class (i.e., coding SNVs [*CS*], coding indels [*CI*], intronic SNVs [*IS*], intronic indels [*II*]) and within the same functionality score and minor allele frequency thresholds. Step 4, Cauchy-combination of significance values achieved using different functionality prediction scores, is not pictured here (Methods).

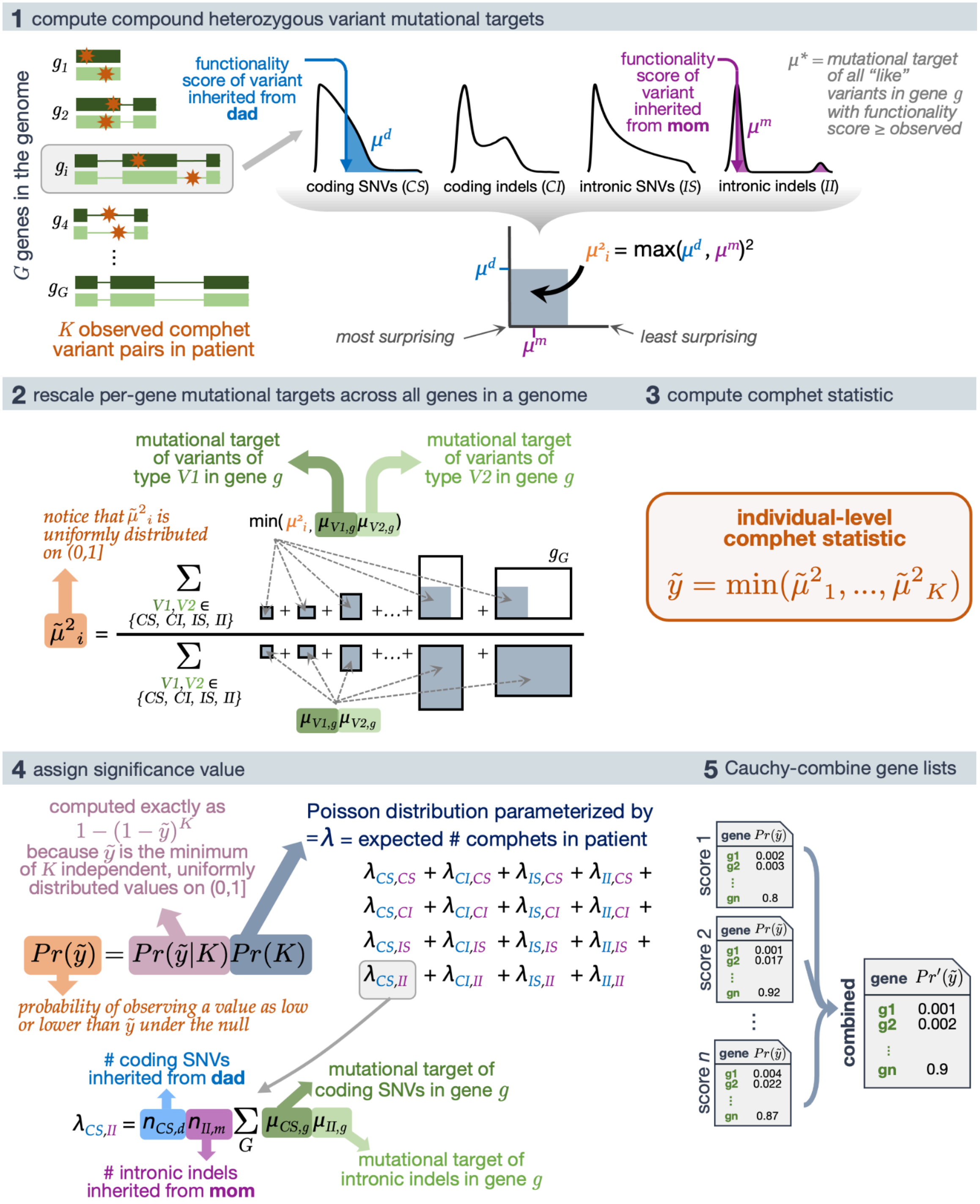

**Figure S12. Schematic of "goodness-of-fit" test for individual-level compound heterozygous variants**

Illustration of compound heterozygous (comphet) variants observed in an individual and simplified graphics depicting the calculation of a comphet mutational target, the rescaling of that mutational target with respect to all *G* genes in a genome to compute an individual-level statistic, and the significance calculation for that individual-level comphet statistic. "Like" variants refer to those of the same variant class (i.e., coding SNVs [*CS*], coding indels [*CI*], intronic SNVs [*IS*], intronic indels [*II*]) and within the same functionality score and minor allele frequency thresholds.

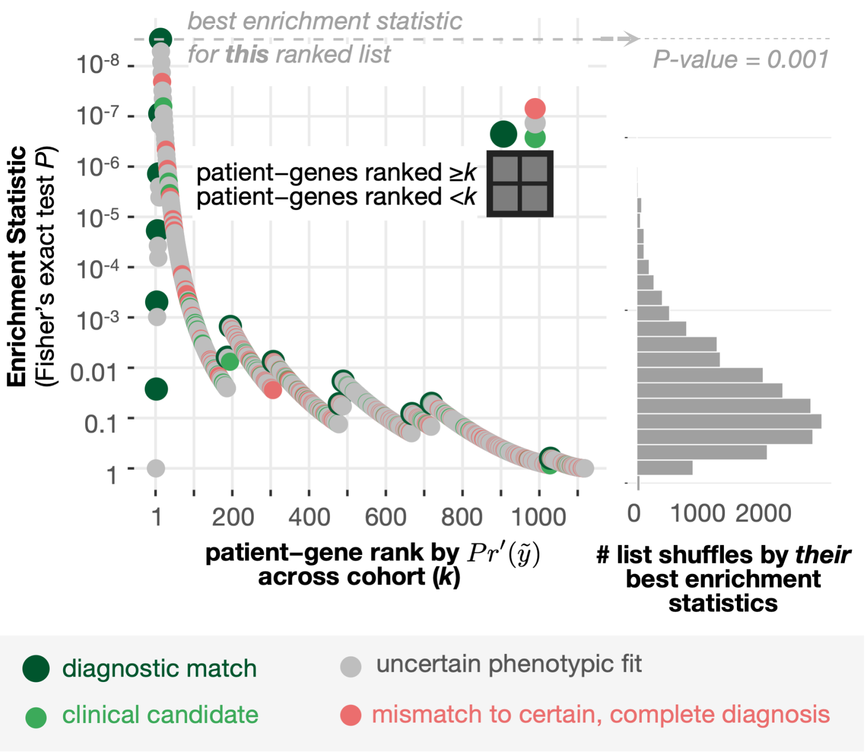

**Figure S13. Enrichment of correct diagnoses in RaMeDiES-IND's patient–gene ranking**
Fisher's exact test *P* values indicate an enrichment of correct diagnoses (dark green circles) among top-ranked genes in a run of RaMeDiES-IND using AlphaMissense, PrimateAI-3D, CADD, and REVEL (left). The 2x2 grid indicates how patient–gene values were counted at each position *k* in the ranked list to compute Fisher's exact test *P*. This enrichment is significant based on the minimum Fisher's exact test *P*s achieved across 10,000 random shuffles of the gene list (right).

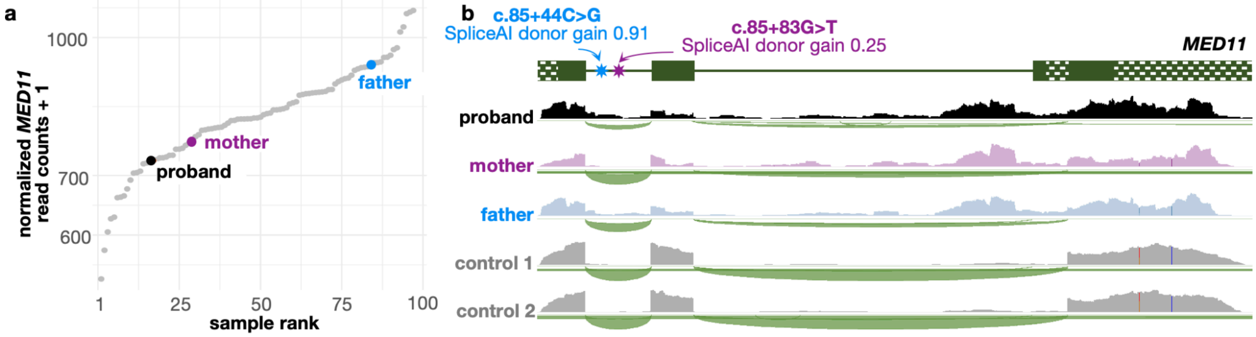

**Figure S14. *MED11* transcriptome sequencing.**
**(a)** Overall gene expression of *MED11* in blood samples from affected proband (black), mother (purple) and father (blue) are within the normal range of expression when compared to 50 sequenced control samples (gray) and would not have been flagged as differentially expressed.
**(b)** RNA-sequencing read depth over the full *MED11* gene region for proband, mother, father, and two control samples.

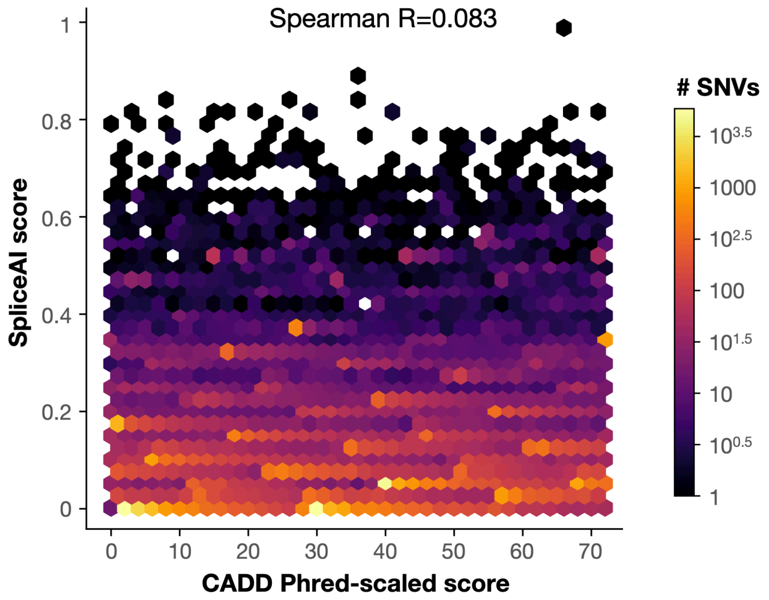

**Figure S15. Correlation between CADD and SpliceAI scores in intronic regions.**
We assessed the correlation between precomputed *in silico* CADD deleteriousness and SpliceAI functionality scores within all intronic regions between protein-coding exons on chromosome 21. We observed poor correlation between these scores, even though CADD uses SpliceAI as an input feature, supporting our distinction between these score types in introns.

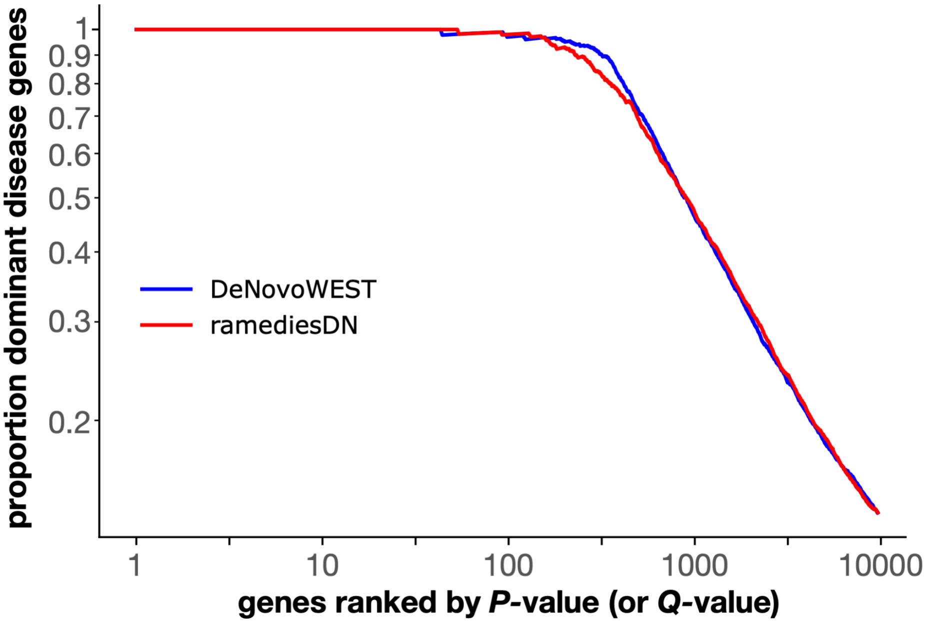

**Figure S16. Comparison of DeNovoWEST and RaMeDiES-DN (CADD only)**
Curves depict the ability of the two *de novo* recurrence methods to recover known autosomal dominant disease genes (reported in OMIM) from the DDD+GeneDx+RUMC dataset. Around 4000 putative *de novos* from the DDD+GeneDx+RUMC were removed because they were on chromosome X, were not in standard VCF format (i.e., non-normalized or left-aligned indels), or could not be mapped to GRCh38. Both axes are log-scaled for visual clarity. A version of RaMeDiES-DN using only CADD scores in exonic regions was used here. DeNovoWEST genes are ranked by *P*-value, and RaMeDiES-DN genes are ranked by *Q*-value.

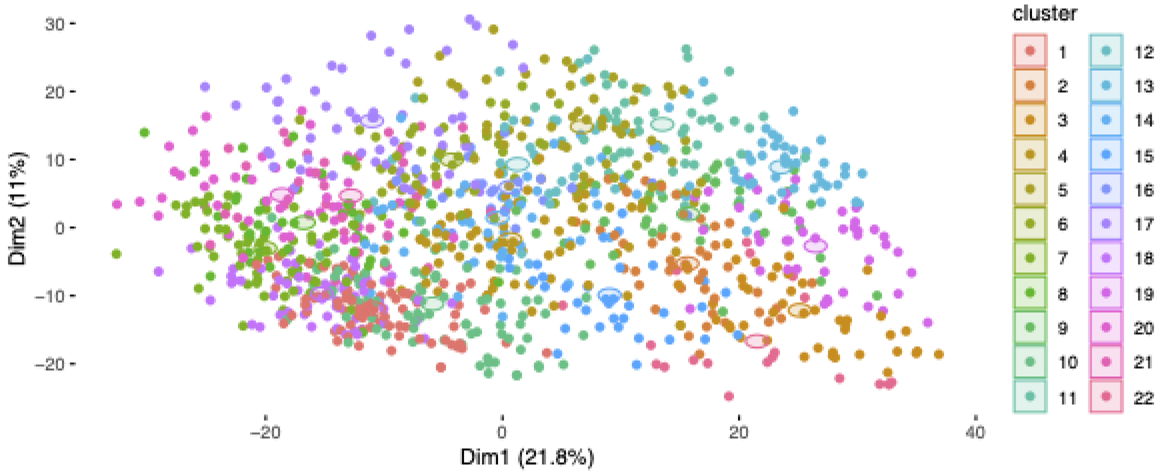

**Figure S17. UDN patients do not fall into well-defined phenotypic clusters.**
UDN patients were clustered using K-medoids clustering based on their pairwise phenotypic similarity score; 22 clusters were selected for visual clarity. Patients separated by pairwise similarity and colored by cluster membership are visualized with a Uniform Manifold Approximation and Projection (UMAP) to show that patients do not easily separate into equally-sized and well-defined clusters.

### Tables

##### Table S1. Candidate vs. Decoy Genes and Final Clinical Scores

Clinical team applying protocol to assign final clinical scores (descending values from most to least compelling) was blinded to gene labels (i.e., decoy or candidate) and *in silico* variant pathogenicity predictions.

| gene | patient phenotype | score | label |
|---|---|---|---|
| AHR | global brain atrophy, history of pituitary gland inflammation (causing hormone deficiency), hypothalamus-related obesity, reduced bone density, history lung blood clots (treated with Eliquis) | 4 | decoy |
| MYO18B | suspected neurodegeneration with brain iron accumulation | 5 | decoy |
| LTBP4 | chronic face/lip swelling, soft tissue thickening (on MRI), inflamed lips with lumps or nodules, tongue with cracks or grooves, high blood pressure, heart mitral valve prolapse | 4 | decoy |
| LRPPRC | adult-onset sudden blurry vision, history of demyelinating disease (white matter, gray matter, cerebellum), double vision, lack of coordination and balance (ataxia), tremor, difficulty swallowing (dysphagia), peripheral sensory neuropathy, motor speech disorder (dysarthria), heart atrial fibrillation, lower extremity weakness (proximal > distal and left > right) | 4 | decoy |
| PNPT1 | connective tissue problems (cutis laxa), pouch-like protrusions (diverticula) in gastrointestinal tract from esophagus to colon, diverticula | 3 | decoy |

| | | | |
|---|---|---|---|
| | in bladder, left abdominal hernia (repaired 3x), weakened heart muscles not due to blocked arteries (non-ischemic cardiomyopathy), thickened heart mitral valve with calcium deposits, normal heart aortic valve | | |
| DYSF | severely high blood pressure (refractory hypertension), vision (ophthalmologic) disturbances, inability to eat solid foods, asymmetric tremor, chronic rash, connective tissue problems (cutis laxa), skin thickening (hyperkeratosis), mild heart aortic dilation, narrowing of renal arteries, low level of white blood cells (lymphopenia) | 5 | decoy |
| OTOGL | autoimmune skin condition (discoid lupus), rheumatoid arthritis, weakened heart with cardioverter-defibrillator device implanted, cerebellar stroke with severe tremor, anemia, low platelets, abnormal protrusion of shoulder blades, scoliosis requiring surgical intervention, progressive muscle loss (cachexia), severe muscle pain, fatigue with lack of endurance | 4 | decoy |
| CEP164 | adolescent-onset stiffness, weakness and cramping after exercise lasting up to a week accompanied by dark urine | 5 | decoy |
| CLCN1 | reduced muscle tone, failure to thrive (improved with gastrostomy tube), gastrointestinal issues (impaired colon movement, extra tissue pushing anus forward, misaligned intestines), lung issues (high blood pressure in arteries leading to lungs, widening of bronchial airways, abnormalities in interstitial tissue) | 3 | decoy |
| PRG4 | enlarged lymph nodes in abdomen with small clusters of immune cells (non-caseating granulomas), elevated white blood cells (eosinophils) in peripheral blood, inflammation around portal veins in the liver | 5 | decoy |
| KCNH8 | feet turning inward (progressively worsening), knee contractures, normal upper extremities, streak-like decreased skin pigmentation with rough patches on left side of neck and arm, delayed muscle relaxation after contraction, decreased reflexes | 2 | candidate |
| NEK8 | refractory epilepsy, movement disorder (dystonia, choreoathetosis), cortical visual impairment, intellectual disability, concern for mitochondrial dysfunction, adrenal insufficiency | 5 | candidate |
| FAN1 | Turner Syndrome, high blood pressure with unknown cause, generalized epilepsy, central auditory processing disorder, intellectual disability, concern for regression. | 3 | candidate |
| ITPR3 | global developmental delay, impaired coordination and balance (ataxia), abnormally large head, autism, cerebral palsy | 2 | candidate |
| SPTBN5 | global developmental delay, absent speech, low muscle tone, failure to thrive, significantly reduced reflexes | 2 | candidate |
| PPP6R1 | severe adult-onset sensory polyneuropathy beginning with numbness and tingling in feet and hands, dizziness with vertigo, diminished and distorted sense of taste | 3 | candidate |
| UNC5C | slowly progressive spinal cord dysfunction (myelopathy), compressed nerve roots (radiculopathy), peripheral nerve damage (neuropathy) | 1 | candidate |
| TPP2 | elevated inflammatory markers, elevated IgM and IgG, neuropathy, chronic pain, cardiomyopathy, cognitive impairment, dysmorphic features, short stature, excessive growth of coarse, dark hair in patterns typical in males (hirsutism), abnormally small red blood cells (microcytic anemia) | 2 | candidate |

| | | | |
|---|---|---|---|
| WDR96 | impaired coordination and balance (ataxia), speech disorder due to weak speech production muscles (dysarthria), difficulty swallowing (dysphagia), progressive cerebellar volume loss shown on MRI | 5 | candidate |
| COL3A1 | adult-onset oral and genital ulcers, extensive skin rash with itchy, small reddish-brown bumps lasting over a year | 4 | candidate |

**Table S2.** ***De novo*** **recurrence findings using AI pathogenicity predictors for missense variants**

We computed our *de novo* recurrence statistic (RaMeDiES-DN) across confident *de novo* variants from 872 affected individuals as described in Methods. We applied two AI-based pathogenicity predictors, AlphaMissense and PrimateAI-3D, which score missense variants only. Columns (from left to right) correspond to:

1. gene name (HGNC ID),
2. Ensembl gene ID,
3. Cauchy-combined P-value (uncorrected for multiple hypotheses),
4. patient UDN diagnosis status (correct gene match, UDN clinician-derived candidate, *incorrect* gene match with an alternate diagnosis listed, or blank),
5. unadjusted patient–gene HPO term similarity score (computed with Phrank),
6. whether the gene is a known developmental disorder gene[3]
7. patient sex,
8. variant consequence,
9. variant functionality score(s),
10. a comment column reminding users that a patient–gene similarity score may be 0 despite a correct diagnosis for the patient if HPO annotations for the patient or gene were limited,
11. the GeneBayes constraint-weighted Q-value (uncorrected for multiple hypotheses),
12. whether the gene's P-value passed the Bonferroni significance threshold,
13. whether the gene's Q-value passed the equivalent threshold,
14. whether the gene's Q-value is within FDR 5%, and
15. whether the gene's Q-value is within FDR 10%.

**Table S3.** ***De novo*** **recurrence findings using four pathogenicity predictors for exonic variants**

We computed our *de novo* recurrence statistic (RaMeDiES-DN) across confident *de novo* variants from 872 affected individuals as described in Methods. We applied four pathogenicity predictors, three of which score missense variants only (i.e., AlphaMissense, PrimateAI-3D, and REVEL) and one of which additionally scores nonsense and indel variants (CADD). Columns (from left to right) correspond to:

1. gene name (HGNC ID),
2. Ensembl gene ID,

3. Cauchy-combined P-value (uncorrected for multiple hypotheses),
4. patient UDN diagnosis status (correct gene match, UDN clinician-derived candidate, *incorrect* gene match with an alternate diagnosis listed, or blank),
5. unadjusted patient–gene HPO term similarity score (computed with Phrank),
6. whether the gene is a known developmental disorder gene[3]
7. patient sex,
8. variant consequence,
9. variant functionality score(s),
10. a comment column reminding users that a patient–gene similarity score may be 0 despite a correct diagnosis for the patient if HPO annotations for the patient or gene were limited,
11. the GeneBayes constraint-weighted Q-value (uncorrected for multiple hypotheses),
12. whether the gene's P-value passed the Bonferroni significance threshold,
13. whether the gene's Q-value passed the equivalent threshold,
14. whether the gene's Q-value is within FDR 5%, and
15. whether the gene's Q-value is within FDR 10%.

**Table S4. Individual-level compound heterozygous findings**

We computed our individual-level compound heterozygous statistic across rare inherited variants from 854 affected individuals as described in Methods. Columns (from left to right) correspond to:

1. Cauchy-combined P-value (uncorrected for multiple hypotheses),
2. gene name (HGNC ID),
3. Ensembl gene ID,
4. patient UDN diagnosis status (correct gene match, UDN clinician-derived candidate, *incorrect* gene match with an alternate diagnosis listed, or blank),
5. unadjusted patient–gene HPO term similarity score (computed with Phrank),
6. patient sex,
7. maternal variant consequence (if known),
8. maternal variant functionality scores (if known),
9. paternal variant consequence (if known),
10. paternal variant functionality scores (if known) .

**Table S5. Candidate genes per patient cluster**

Patients were clustered into phenotypically-similar subgroups, and then compelling genes were selected per patient in each cluster as described in Methods. The set of genes per patient cluster was used as a query for gene set enrichment analysis. Columns (from left to right) correspond to: pathway ID ("HC" indicates hierarchical clustering), gene name (HGNC ID), gene type (i.e., "denovo" indicates a compelling *de novo* variant was present, "comphet" indicates a compelling compound heterozygous variant was present, and "augmented" indicates a known diagnosis that was established through the UDN process), patient diagnosis status, and unadjusted patient–gene HPO term similarity score (computed with Phrank). Note that some correct diagnoses may have a patient–gene phenotype similarity score of zero if either the patient or the gene did not have any annotated HPO terms.

**Table S6. All enriched biological pathways across patient clusters**
We performed gene set enrichment analysis using Reactome and KEGG biological pathways as described in Methods. The query genes selected per patient cluster include known diagnoses achieved through UDN analyses as well as computationally-derived *de novo* and compound heterozygous candidates, as described in Methods and listed in Supplementary Table S5. Columns (from left to right) correspond to: pathway ID (asterisk indicates a patient–gene pair that also appears earlier in the list), pathway name, pathway size, GSEA adjusted *p*-value, cluster ID and effective gene set query size, gene name (HGNC ID), Ensembl gene ID, patient diagnosis status, unadjusted patient–gene HPO term similarity score (computed with Phrank), patient sex, variant consequence (if known), variant functionality scores (if known). Note that the effective gene set query size can be different for the same cluster ID between Reactome and KEGG enriched pathways. The original gene set query for each patient cluster is intersected with the set of annotated genes across all Reactome or all KEGG pathways respectively; some genes may only be present in one of the two data sources.

### Notes

#### Note S1. Subdividing neurological symptom category

To identify subgroups within the largest "neurological" primary symptom category, we used the all-against-all pairwise phenotypic similarity scores previously computed (Methods) to cluster the subset of neurological patients into three equivalently-sized groups (using Ward-Linkage hierarchical clustering from the R cluster package). We created a text corpus for each cluster from the concatenated list of patients' HPO terms within each cluster using the R tm package and visualized these text corpuses as comparative word clouds using the R quanteda package. Subgroup names were determined based on manual inspection and comparison of the word clouds.

#### Note S2. Clinical evaluation protocol

In all of the following sections, an asterisk (*) indicates fields that were extracted computationally in our implementation.

**Preparation: Collecting information prior to evaluating diagnostic fit of candidate**
Compile relevant information in your database of choice; Microsoft Excel spreadsheets and REDCap databases have both been used effectively in following this protocol.

***Information collected about the affected patient:***

1. Patient ID*
2. Family ID*
3. Clinical site of evaluation*
4. Sequencing lab where genome sequencing was completed*
5. RNA-Seq available? If so: sequence ID and tissue*
6. Known diagnosis available? If so: gene name, diagnosis certainty level, diagnosis completeness (complete or partial)*
7. Other candidate genes available? If so: gene name*
8. Heritage/demography*
9. Age at symptom onset*
10. Age at application to UDN and age at clinical evaluation*
11. Sex*
12. Affected family members? If so: their patient IDs*
13. Human Phenotype Ontology (HPO) codes and terms*
14. Abstracted case review of patient presentation (based on Application Letter, Wrap Up Form/Letter, Photography and Imaging Reports, see Methods)*

15. Clinician-determined primary symptom category*
16. Affected tissue(s) - manually determined, and used to assess relevant tissue-specific expression later

***Information collected about each variant:***

1. Variant classification (exonic or intronic)*
2. Genome build*
3. Variant location = chromosome, position, reference DNA allele(s), alternate DNA allele(s)*
4. Variant read depth across trio*
5. Variant impact from Ensembl's Variant Effect Predictor*
6. Variant transcript = transcript name and HGVS cDNA impact*
7. Variant protein = protein name and HGVS protein impact*
8. CADD score (exonic and intronic)*
9. SpliceAI score (exonic and intronic)*
10. Confirmed in IGV? (i.e., present on a clinical sequencing report or in a research variant table)

***Tip for planning for literature review***

Before beginning, note that there are **three** separate fields where PubMed IDs and brief take-home summaries should be recorded; in practice the PubMed search happens *once* and you will record relevant information in the different corresponding fields:

(1) Articles associating gene to known human diseases (matching patient phenotype or not)[†]
(2) Articles about protein molecular function[†]
(3) Articles about experiments (engineered cell lines or animal models) relating gene variants to phenotype changes[‡]

[†]These articles are sometimes curated in OMIM, but thorough review of PubMed is necessary for the most up-to-date evidence review.
[‡]This information is sometimes found in MGI (Mouse Genome Informatics); record PubMed IDs where possible.

**Evaluation of variant–patient phenotypic fit by category**

External databases are consulted to compile information that will then be used to evaluate the diagnostic fit of the candidate variant(s) for the patient. Here we provide example databases used by our clinical team to execute this protocol, but newer databases not explicitly listed here that compile the same information can be used as well. There are six categories of information to be collected for review.

***Known disease associations***

1. Notes from Clinical Genome Resource (ClinGen[4]) diseases & expert panel, Gene Curation Coalition (GenCC[5]), and Genomics England PanelApp[6]
2. PubMed articles (PMID and brief take-home summary) relating gene to phenotype or gene to patient presentation
    a. **Note:** keep track of supporting as well as contradictory evidence
3. Manually-written summary of interesting information from the above step 2
4. OMIM disease associations and their IDs*
5. Published reports of disease/gene associations from OMIM[7] (obtained via manual review of OMIM pages)
6. Associated traits in gene from GWAS catalog[8] / linkage associations
    a. Sometimes there are hundreds of traits, so we take the top/most relevant 15-20
7. GeneMatcher (MatchMaker Exchange)
8. Phrank score relating gene to patient's phenotype terms (low scores are unreliable but high scores are interesting)*

| *Do any known disease associations overlap with proband's key phenotypic features?* | |
|---|---|
| 1 | ClinGen curated disease association with correct inheritance mode (i.e., AD for *de novos* or AR for compound heterozygous variants) with strong phenotypic overlap with patient's primary phenotype(s)/pathophysiology |
| 2 | Published report or OMIM annotation with correct inheritance mode supporting disease monogenic association with strong phenotypic overlap |
| 3 | Any published report/ClinGen curation with mild-moderate phenotypic overlap (regardless of inheritance mode) |
| 4 | Published report/OMIM annotation supporting disease polygenic/GWAS association with phenotypic overlap |
| 5 | No evidence supporting any gene/disease association |
| 6 | Known gene-disease association with minimal phenotypic overlap (regardless of inheritance mode) |
| 7 | Known disease association with correct inheritance mode with no phenotypic overlap |

***Model organism studies***

1. Mouse Genome Informatics (MGI)[9] > search for gene name (and click symbol in results table) > Mutations, Alleles and Phenotypes section > review summary at bottom
2. International Mouse Phenotyping Consortium (IMPC)[10] can also be consulted, similar to information available in MGI.
3. Review PubMed articles about experimental evidence (engineered cell lines or animal models) relating variants to phenotypes.
4. Models and Rescue score, max 4, (based on information from MGI, IMPC, and PubMed):[11]
    a. +2 = Animal model rescue: Was the variant introduced and did it result in a matching phenotype in the animal, and then was the variant reversed/blocked via a CRISPR/Cas9 or silencing RNA intervention that reversed the phenotype? (1 point for cell culture model or engineered equivalent)
    b. +1 = Functional alteration in cell model (0.5 engineered, 1 for cell line from affected individual)

| *Does tissue-level expression in model organisms or null models suggest correlation with proband phenotypes?* | |
|---|---|
| 1 | Null mouse shows phenotypic overlap with proband |
| 2 | Other model organism data demonstrate overlap with proband phenotypic features |
| 3 | Expression in other organisms overlaps with key phenotypic features/tissues of interest |
| 4 | Expression in other organisms overlaps mildly with key phenotypic features/tissues of interest |
| 5 | Expression in other organisms does not overlap with key phenotypic features |
| 6 | No model organism data available |

***Expression of gene in phenotypically-relevant tissues***

1. Check expression in the Allen Brain Institute's Brain Map[12]
2. The Genotype-Tissue Expression (GTEx) Portal[13] – search for gene, scroll down to "Bulk tissue gene expression" to see relevant tissues

| Does available tissue-level expression data align with proband's key phenotypic features? | |
|---|---|
| 1 | Multiple datasets show tissue expression / phenotypic feature overlap |
| 2 | At least one dataset shows tissue expression / phenotypic features overlap |
| 3 | Available data shows minimal tissue expression of gene |
| 4 | Datasets show expression in tissues other than those related to proband's phenotypic features |

***Protein molecular function***

1. Entrez gene summary (reviewed via GeneCards[13,14])
2. UniprotKB/SwissProt protein summary (reviewed via GeneCards[15])
3. Published reports of gene/phenotype associations from OMIM[7,15]

| *Does the protein function relate to the potential model of molecular pathophysiology?* | |
|---|---|
| 1 | Well-annotated gene product function with strong association with patient's primary phenotypic feature(s)/pathophysiology |
| 2 | Published report/OMIM annotation supporting monogenic disease association with mild phenotypic overlap |
| 3 | Minimal or no gene product function annotation |
| 4 | Well-annotated gene product function with no obvious association with patient's primary phenotype(s)/pathophysiology |

***Gene evolutionary constraint***

1. Loss-of-function (nonsense, splice acceptor, splice donor) scores from gnomAD*[16]
    a. pLI (ranges 0-1) where 1 means *intolerant* to LoF variants
    b. Observed/expected ratio (ranges 0-1) and Poisson-derived confidence interval; where 0 means *intolerant* to LoF variants; note that the upper bound of the Poisson confidence interval is referred to as "LOEUF" for "loss-of-function observed/expected upper bound fraction"
2. Missense scores from gnomAD* browser: Z-score (positive value = more constraint) and o/e ratio and confidence interval (where upper bound of the Poisson-derived confidence interval is called MOEUF) from gnomAD* browser

    a. Missense Z-score > 3.09 is more likely to be intolerant of missense changes
3. UCSC PhastCons100Way score (higher means more conserved) and manual notes from visual inspection (e.g., "conserved in all but zebrafish")
    a. PhyloP ranges from -14 to 3, with higher values indicating more conserved
    b. GERP++ is recommended for use in ClinGen (again, higher is more conserved).

*In our implementation, gnomAD v2.1.1 was used for exomes and v3.1.2 for genomes, but any subsequent gnomAD releases should be applicable for these steps as well.

| *Is there evidence of evolutionary constraint at the gene level?* | |
|---|---|
| 1 | Constrained for LOF variation (LOEUF<0.35) and constrained for missense variation (z-mis>+2) |
| 2 | Constrained for LOF variation (LOEUF<0.35) |
| 3 | Constrained for missense variation (z-mis>+2) |
| 5 | Moderate evidence for constraint (LOEUF<0.5*** or z-mis>+1) |
| 4 | Minimal to no evidence of constraint |

***Variant frequency, pathogenicity, and position***

**Note:** This section is only completed *if* the gene seems to be a good phenotypic match based on the previous sections.

1. Maximum allele frequency (gnomAD* population maximum)*
2. GnomAD* population group corresponding to popmax
3. Number of homozygotes (for compound heterozygous candidates) and number of heterozygotes (for *de novo* dominant candidates) (gnomAD*)
4. Computational pathogenicity predictors (# deleterious / # tolerated), reviewed from VarSome[17,18]
5. ClinVar categorization and star category*
6. Proximity to other ClinVar variants (from visual inspection in DECIPHER[19] database Protein/ Genomic track view)
7. Variant location relative to protein functional domains (from visual inspection in DECIPHER and UniProt)
8. Missense depleted area (from visual inspection in DECIPHER gnomAD* Missense track)

    a. DECIPHER > search for gene > Protein/Genomic tab > scroll down to tracks > gnomAD* missense track
9. Tissue expression of exon containing variant (gnomAD*)
    a. From the gnomAD* browser, search for the gene name > copy the HGVS protein position into the field under “gnomAD* variants” > hover over location where (coding) variant is then scroll up through the tissues > hover over exon in each tissue and note the “pext” score (ranges 0=low to 1=high)

*In our implementation, gnomAD v2.1.1 was used for exomes and v3.1.2 for genomes, but any subsequent gnomAD releases should be applicable for these steps as well.

| *Does the variant seem pathogenic?* | |
|---|---|
| 1 | Known/likely pathogenic variant |
| 2 | VUS - suspect likely pathogenic |
| 3 | VUS |
| 4 | VUS - suspect likely benign |
| 5 | Known benign variant |

**Summarizing information for final phenotypic fit score**
Once information has been filled in and scores assigned for the intermediate steps, at least two clinicians will meet to go over the scores. We recommend keeping track of per-clinician notes and the group consensus.

1. Notes from individual clinicians (if any)
2. Evidence supporting association
3. Evidence against association
4. Reviewed as a group? (yes/no)
5. Notes from group meeting (if any)

| *Summary Classification* | |
|---|---|
| 1 | Known disease association with strong phenotypic overlap with proband |
| 2 | Multiple lines of supporting evidence for association with proband phenotype |
| 5 | Multiple weak lines of evidence for association with proband phenotype |

| | |
|---|---|
| 3 | One line of supporting evidence for association with proband phenotype or conflicting evidence |
| 4 | No evidence for gene-phenotype association |

#### Note S3. Compound heterozygous mutational target is uniformly distributed

Recall that we define the mutational target of a compound heterozygous variant configuration as

$$\mu_{g,v_M,v_D} = \max(\mu_{g,v_M}, \mu_{g,v_D})^2.$$

We will show here that, by this definition, $\mu_{g,v_M,v_D}$ is uniformly distributed. Let $\mu_1$ and $\mu_2$ be dummy variables corresponding to some maternal and paternal variant mutational targets in a compound heterozygous configuration in gene $g$, and let $\mu_g$ be the mutational target of gene $g$.

$$Pr(\mu_{g,v_M,v_D}) = Pr(\mu_{g,v_M,v_D} \geq \max(\mu_1, \mu_2)^2)$$

$$= Pr(\mu_{g,v_M,v_D} \geq \max(\mu_1^2, \mu_2^2))$$

$$= Pr\left(\frac{\mu_{g,v_M,v_D}}{\mu_g^2} \geq \max\left(\frac{\mu_1^2}{\mu_g^2}, \frac{\mu_2^2}{\mu_g^2}\right)\right)$$

Suppose $\max(\mu_1, \mu_2) = \mu_1$.

$$= Pr\left(\frac{\mu_{g,v_M,v_D}}{\mu_g^2} \geq \frac{\mu_1^2}{\mu_g^2}\right) Pr\left(\frac{\mu_{g,v_M,v_D}}{\mu_g^2} \geq \frac{\mu_1^2}{\mu_g^2}\right)$$

$$= Pr\left(\frac{\mu_{g,v_M,v_D}}{\mu_g^2} \geq \frac{\mu_1^2}{\mu_g^2}\right)^2$$

$$= Pr\left(\sqrt{\frac{\mu_{g,v_M,v_D}}{\mu_g^2}} \geq \frac{\mu_1}{\mu_g}\right)^2$$

Let $x = \sqrt{\mu_{g,v_M,v_D}/\mu_g^2}$. Recall that, by construction, variant mutational targets $\mu_{g,v_M}$ and

$\mu_{g,v_D}$ are both $\leq \mu_g$ and so $\mu_{g,v_M,v_D} \leq \mu_g^2$, and $\mu_{g,v_M,v_D}/\mu_g^2$ and its square root must range from 0 to 1 (Equation 1, Methods). So, $x \in [0,1]$.

$$= Pr\left(x \geq \frac{\mu_1}{\mu_g}\right)^2$$

We have already established that $\mu_1/\mu_g$ is uniformly distributed on [0, 1] . Since the cumulative distribution function of a uniformly distributed variable on $[a,b]$ is $(x-a)/(b-a)$ for $x \in [a,b]$ and therefore just $x$ when $x \in [0,1]$, we simplify the above expression as

$$= x^2 = \left(\sqrt{\frac{\mu_{g,v_M,v_D}}{\mu_g^2}}\right)^2 = \frac{\mu_{g,v_M,v_D}}{\mu_g^2}$$

Since we have shown that $Pr(\mu_{g,v_M,v_D}) = \mu_{g,v_M,v_D}/\mu_g^2$, by the same logic applied above, it follows that

$$\frac{\mu_{g,v_M,v_D}}{\mu_g^2} \sim \mathcal{U}_{[0,1]}$$

#### Note S4. Independence assumption is violated in homozygous recessive case

Our individual-level compound heterozygous configuration statistic $\tilde{y}^c$ (Equation 5, Methods) and the expected number of genes harboring compound heterozygous configurations per individual genome $\tilde{\lambda}^c$ (Equation 6, Methods) both assume that the two inherited variants comprising a compound heterozygous configuration arose as a result of independent mutations. In the case of homozygous recessive biallelic variants, however, where the exact same variant is inherited from each parent, this independence assumption is violated because in most cases a single mutational event resulted in the presence of a variant in both parents. Given this violation, we directly and stringently exclude individuals with evidence of consanguinity (i.e., requiring parental relatedness <0.15 and maximum IBD length < 3Mb) rather than apply a simple test to filter individuals with relatively high recessive burden.

#### Note S5. Undiagnosed Diseases Network Consortium Members

Maria T. Acosta[1], David R. Adams[1], Ben Afzali[1], Ali Al-Beshri[2], Eric Allenspach[3], Aimee Allworth[3], Raquel L. Alvarez[4], Justin Alvey[5], Ashley Andrews[5], Euan A. Ashley[6], Carlos A.

Bacino[7], Guney Bademci[8], Ashok Balasubramanyam[7], Dustin Baldridge[9], Jim Bale[5], Michael Bamshad[3], Deborah Barbouth[8], Pinar Bayrak-Toydemir[5], Anita Beck[3], Alan H. Beggs[10], Edward Behrens[11], Gill Bejerano[4], Hugo J. Bellen[12], Jimmy Bennett[3], Jonathan A. Bernstein[4], Gerard T. Berry[10], Anna Bican[13], Stephanie Bivona[8], Elizabeth Blue[3], John Bohnsack[5], Devon Bonner[4], Nicholas Borja[8], Lorenzo Botto[5], Lauren C. Briere[10], Elizabeth A. Burke[1], Lindsay C. Burrage[7], Manish J. Butte[14], Peter Byers[3], William E. Byrd[2], Kaitlin Callaway[2], John Carey[5], George Carvalho[14], Thomas Cassini[13], Sirisak Chanprasert[3], Hsiao-Tuan Chao[7], Ivan Chinn[7], Gary D. Clark[7], Terra R. Coakley[4], Laurel A. Cobban[10], Joy D. Cogan[13], Matthew Coggins[10], F. Sessions Cole[15], Brian Corner[13], Rosario I. Corona[14], William J. Craigen[7], Andrew B. Crouse[2], Vishnu Cuddapah[11], Precilla D'Souza[1], Hongzheng Dai[7], Kahlen Darr[16], Surendra Dasari[16], Joie Davis[1], Margaret Delgado[1], Esteban C. Dell'Angelica[14], Katrina Dipple[3], Daniel Doherty[3], Naghmeh Dorrani[14], Jessica Douglas[10], Emilie D. Douine[14], Dawn Earl[3], Lisa T. Emrick[7], Christine M. Eng[17], Cecilia Esteves[18], Kimberly Ezell[13], Elizabeth L. Fieg[10], Paul G. Fisher[4], Brent L. Fogel[14], Jiayu Fu[1], William A. Gahl[1], Rebecca Ganetzky[11], Emily Glanton[18], Ian Glass[3], Page C. Goddard[4], Joanna M. Gonzalez[8], Andrea Gropman[1], Meghan C. Halley[4], Rizwan Hamid[13], Neil Hanchard[1], Kelly Hassey[11], Nichole Hayes[9], Frances High[10], Anne Hing[3], Fuki M. Hisama[3], Ingrid A. Holm[10], Jason Hom[4], Martha Horike-Pyne[3], Alden Huang[14], Yan Huang[1], Anna Hurst[2], Wendy Introne[1], Gail P. Jarvik[3], Suman Jayadev[3], Orpa Jean-Marie[1], Vaidehi Jobanputra[19], Oguz Kanca[12], Yigit Karasozen[14], Shamika Ketkar[7], Dana Kiley[9], Gonench Kilich[11], Eric Klee[16], Shilpa N. Kobren[18], Isaac S. Kohane[18], Jennefer N. Kohler[4], Bruce Korf[2], Susan Korrick[10], Deborah Krakow[14], Elijah Kravets[4], Seema R. Lalani[7], Christina Lam[3], Brendan C. Lanpher[16], Ian R. Lanza[16], Kumarie Latchman[8], Kimberly LeBlanc[18], Brendan H. Lee[7], Kathleen A. Leppig[3], Richard A. Lewis[7], Pengfei Liu[17], Nicola Longo[5], Joseph Loscalzo[10], Richard L. Maas[10], Calum A. MacRae[10], Ellen F. Macnamara[1], Valerie V. Maduro1, AudreyStephannie C. Maghiro[18], Rachel Mahoney[18], May Christine V. Malicdan[1], Rong Mao[5], Ronit Marom[7], Gabor Marth[20], Beth A. Martin[4], Martin G. Martin[14], Julian A. Martínez-Agosto[14], Shruti Marwaha[4], Allyn McConkie-Rosell[21], Ashley McMinn[13], Matthew Might[2], Mohamad Mikati[21], Danny Miller[3], Ghayda Mirzaa[3], Breanna Mitchell[16], Paolo Moretti[5], Marie Morimoto[1], John J. Mulvihill[1], Lindsay Mulvihill[16], Mariko Nakano-Okuno[2], Stanley F. Nelson[14], Serena Neumann[13], Dargie Nitsuh[3], Donna Novacic[1], Devin Oglesbee[16], James P. Orengo[7], Laura Pace[5], Stephen Pak[9], J. Carl Pallais[10], Neil H. Parker[14], LéShon Peart[8], Leoyklang Petcharet[1], John A. Phillips III[13], Filippo Pinto e Vairo[16], Jennifer E. Posey[7], Lorraine Potocki[7], Barbara N. Pusey Swerdzewski[1], Aaron Quinlan[5], Daniel J. Rader[11], Ramakrishnan Rajagopalan[11], Deepak A. Rao[10], Anna Raper[11], Wendy Raskind[3], Adriana Rebelo[8], Chloe M. Reuter[4], Lynette Rives[13], Lance H. Rodan[10], Martin Rodriguez[2], Jill A. Rosenfeld[7], Elizabeth Rosenthal[3], Francis Rossignol[1], Maura Ruzhnikov[4], Marla Sabaii[1], Jacinda B. Sampson[4], Timothy Schedl[9], Lisa Schimmenti[16], Kelly Schoch[21], Daryl A. Scott[7], Elaine Seto[7], Vandana Shashi[21], Emily Shelkowitz[3], Sam Sheppeard[3], Jimann Shin[9], Edwin K. Silverman[10], Giorgio Sirugo[11], Kathy Sisco[9], Tammi Skelton[2], Cara Skraban[11], Carson A. Smith[8], Kevin S. Smith[4], Lilianna Solnica-Krezel[9], Ben Solomon[1], Rebecca C. Spillmann[21], Andrew Stergachis[3], Joan M.

Stoler[10], Kathleen Sullivan[11], Shirley Sutton[4], David A. Sweetser[10], Virginia Sybert[3], Holly K. Tabor[4], Queenie Tan[16], Arjun Tarakad[7], Herman Taylor[22], Mustafa Tekin[8], Willa Thorson[8], Cynthia J. Tifft[1], Camilo Toro[1], Alyssa A. Tran[7], Rachel A. Ungar[4], Adeline Vanderver[11], Matt Velinder[5], Dave Viskochil5, Tiphanie P. Vogel7, Colleen E. Wahl1, Melissa Walker10, Nicole M. Walley21, Jennifer Wambach[9], Michael F. Wangler[12], Patricia A. Ward[17], Daniel Wegner[9], Monika Weisz Hubshman[7], Mark Wener[3], Tara Wenger[3], Monte Westerfield[23], Matthew T. Wheeler[4], Jordan Whitlock[2], Lynne A. Wolfe[1], Heidi Wood[1], Kim Worley[7], Shinya Yamamoto[12], Stephan Zuchner[8]

**1** National Institutes of Health (NIH) Undiagnosed Diseases Program (UDP) clinical site
**2** University of Alabama at Birmingham, Data Management and Coordinating Center
**3** Pacific Northwest clinical site
**4** Stanford University clinical site
**5** University of Utah clinical site
**6** Stanford University Data Management and Coordinating Center
**7** Baylor College of Medicine clinical site
**8** University of Miami clinical site
**9** Washington University clinical site, Model Organisms Screening Center
**10** Harvard University clinical site
**11** Children's Hospital of Philadelphia and University of Pennsylvania clinical site
**12** Baylor College of Medicine Model Organisms Screening Center
**13** Vanderbilt University clinical site
**14** University of California at Los Angeles clinical site
**15** Washington University Data Management and Coordinating Center
**16** Mayo Clinic
**17** Baylor College of Medicine Sequencing Core
**18** Harvard University Data Management and Coordinating Center
**19** Columbia University clinical site
**20** University of Utah Data Management and Coordinating Center
**21** Duke University clinical site
**22** Morehouse Data Management and Coordinating Center
**23** University of Oregon Model Organisms Screening Center

19. DECIPHER v11.24: Mapping the clinical genome. https://www.deciphergenomics.org/browser.
